# Machine learning based identification of candidate miRNA biomarkers for micro-invasive breast cancer diagnosis

**DOI:** 10.1101/2024.08.16.608025

**Authors:** Jayanta K. Pal, Bhadresh R. Rami

## Abstract

**Abstract:** 

**Purpose:** Early detection of cancer can be done by analyzing miRNA expression patterns. miRNAs play a significant role in biological processes, and they have been identified as one of the major biomarkers in cancer. miRNAs can also be detected in human blood (micro-invasive way of sample collection), which makes the diagnostic procedure much less stressful for the patients. In this article, we emphasize on identification of miRNAs as biomarkers (collected from blood sample) that are associated with breast cancer.

**Methods:** In this investigation we use three breast cancer data sets, obtained from blood samples. A combination of multiple feature selection and classification models is used to classify normal vs cancer samples. In the first stage, the significant miRNAs associated with cancer were selected by (a) classifier assigned weights and (b) feature selection algorithms. In the second stage, we apply multiple classifiers to observe the diagnostic capability of the selected miRNAs for consideration as potential biomarkers.

**Results:** Our miRNA selection stage identified ten miRNAs, which were subsequently analysed using multiple classifiers for their ability to distinguish between normal and cancerous cases. The performance is examined using a 5-fold cross validation technique using multiple measures such as precision, recall, F1-score, and accuracy. We also use a confusion matrix to evaluate the performance of the selected miRNAs. For two out of three datasets, we achieve satisfactory performance in terms of normal vs cancer classification.

**Conclusion:** We observe that high expression levels of miRNA is relatively more important than the sample size, for effective blood-based diagnosis of breast cancer. The novelty of our investigation lies in combining three aspects viz., blood-based breast cancer diagnosis, use of multiple ML based feature selection algorithms to identify the miRNAs associated with breast cancer, evaluating them using various classifiers and the robustness of these ML models in feature selection and classification.

## 1. Introduction

MicroRNAs (miRNAs) are a type of non-coding RNA that do not code for proteins. Rather these RNAs interact with messenger RNAs (mRNA) and inhibit protein translation [1]. Proteins are pivotal to all the cellular functions and one such key function is intra-cellular signaling. Signaling pathways play a key role in regulating cell proliferation, apoptosis. Therefore, abnormalities in miRNAs may result in uncontrolled cell division which is a hallmark of cancer.

It has been identified in earlier studies [2, 3] that deletions / dysregulation of miRNA genes and their expression patterns correlate with the onset of cancer. These were the early set of studies that utilized ML models; where the authors used hierarchical clustering on the expression levels and obtained different clusters based on the tissue locations. They identified that miRNA expression levels can differentiate normal vs cancer samples. Emphasis was on developing algorithms that work based on fold change and p-values leading to the selection of globally deregulated miRNAs in multiple cancers [4]. Blood sample-based miRNA collection and diagnosis of melanoma is also reported [5].

Substantial efforts are ongoing towards cancer detection and miRNA selection using various ML models. Emphasis is given on breast cancer diagnosis [6] using Support Vector Machine (SVM) and then multiple subtypes of the disease are identified by Random Forest (RF). Feature importance is also evaluated using a Shapley value-based method. Even though the investigation demonstrates an effective method for breast cancer diagnosis using ML models, it relied only on two models for classification. Moreover, the usefulness of feature importance determination is not clearly explained. Extensive work has been reported on fifteen cancers and their four stages [7] using a feature selection algorithm (IBCGA) and a classifier (SVM). Similar investigation with early and advanced stage of breast cancer is performed [8]. However, in both cases multiple ML models are not considered during classification.

Usefulness of ML models in cancer diagnosis/detection was established in [9] using several feature selection and classification models. However, the biological significance, such as the benefits of using non-invasive/microinvasive methods of sample collection for diagnostic purposes, are not covered by this study. An investigation [10] on gastric cancer was performed to find out the predictive values of miRNAs in cancer. In this investigation, heatmap analysis was performed to identify the significant miRNAs, and then multiple classifiers (such as SVM, Decision Tree (DT), etc.) were used to assess the classification performance of the identified miRNAs. In this study, a single strategy for miRNA selection was employed, and the importance of non-invasive/micro-invasive sample collection methods was also not considered. Similar studies were performed [11, 12, 13] performed using various feature selection and classification models. Here too, the aspect of non/micro-invasive way of sample collection was not considered. Emphasis was given on determining the effectiveness of the treatment in breast cancer based on the miRNA expression levels [14]. Application of Cox regression-based method to identify the important miRNAs responsible for breast cancer has been reported [15]. In this study, only one model is used for miRNA selection and micro-invasive sampling is also not considered.

It is observed from the above discussed studies that they do not address three key issues simultaneously. The issues are (a) diagnosis using miRNAs collected by non/micro-invasive ways of sample collection, (b) use of multiple feature selection algorithms and (c) use of multiple ML models for classification. The first issue ensures less stressful way of sample collection and the last two helps to make the diagnostic procedure more robust as compared to using a single feature selection algorithm or classifier.

To address the aforesaid issues, we performed our investigations using blood-based sampling and used multiple methods for both the selection and classification of miRNAs. Hence, the novelty of our research article lies in combining three aspects viz., (a) blood sample-based breast cancer diagnosis, (b) use of multiple ML based feature selection algorithms to identify the miRNAs associated with breast cancer, (c) evaluating the performances of these selected miRNAs using various classifiers and the robustness of these ML models in feature selection and classification.

## 2. Materials and Methods

In this investigation we aim to identify features/patterns in miRNA expressions that can help differentiate between normal and cancerous conditions. To meet our objectives, we used three different breast cancer data sets (Table 1) form different countries having both normal and cancer samples. Here the miRNAs were collected from the blood-based samples. We then used seven different methods for selecting miRNAs. Few methods used the feature weights assigned by well-known classifiers such as Extra Tree (ET), SVM, Random Forest (RF). Other methods employed dedicated algorithms for feature selection such as chi-square information measure, MRMR-I [16], MRMR-II [17], FREM [18]. Here MRMR-I refers to an approach where feature selection was performed by maximizing the relevance and minimizing the redundance. The MRMR-II is an upgraded version of the MRMR-I (elaborated in section 2.iv.f). Once miRNAs are selected, we used several ML models (such as extra tree, SVM, RF, k-NN, XGBoost, Naïve Bayes and artificial neural network (ANN) classifiers) for classification of normal vs cancer samples. We analyzed the performance of the selected miRNAs and suggested the best possible pairs of miRNA selection and classification models that help to classify normal vs cancer cases. The steps involved in our investigation are presented in Fig. 1.

**Fig. 1.**
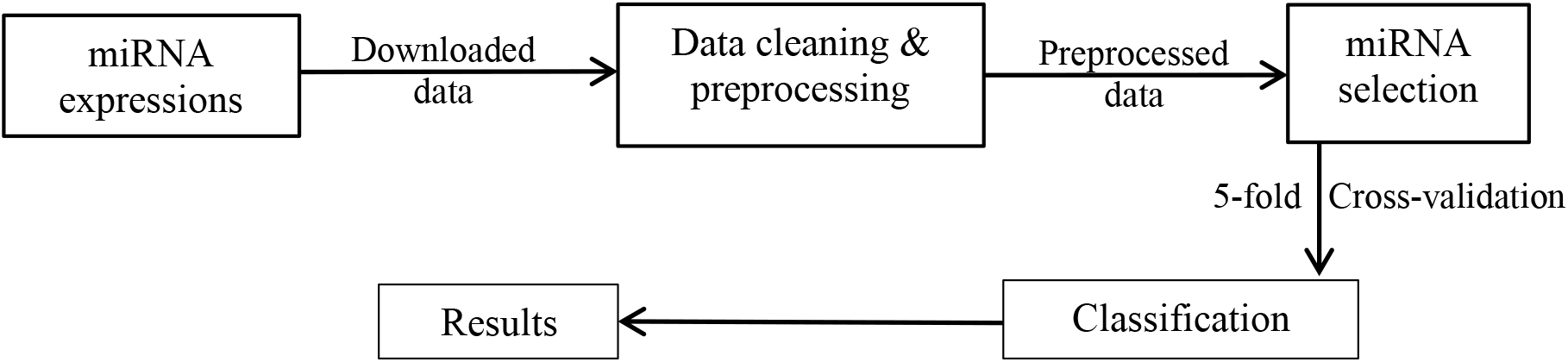
Block diagram.

### i. Data sets

We performed our analysis on publicly available data sets downloaded from Gene Expression Omnibus (GEO), with accession numbers: GSE31309, GSE44281 and GSE87230. The dataset GSE31309 consists of 1100 miRNAs, 57 normal patients & 48 cancer patients. GSE44281 has 1121 miRNAs, 205 patients in each category (i.e., normal and cancer), and GSE87230 contains 1916 miRNAs and 6 patients in each category. The summary of the data sets used in our study is provided in Table 1.

**Table 1:** Data set.

| Study # | Accession | Normal subjects | Cancer patients | miRNAs |
| --- | --- | --- | --- | --- |
| 1 | GSE31309 | 57 | 48 | 1100 |
| 2 | GSE44281 | 205 | 205 | 1121 |
| 3 | GSE87230 | 6 | 6 | 1916 |

### ii. Data cleaning/pre-processing

The miRNA names and the corresponding expression values are stored in different files in the mentioned datasets. Furthermore, miRNAs from species other than humans were also included here, hence, we created a separate file containing only the human miRNA names and their expression levels. Data cleaning is performed as per the steps given below:

a. To create a single file with both entities (i.e., miRNA names and expressions), we first matched the miRNA IDs provided in both files containing miRNA names and expressions. The “Vlookup” function in Microsoft Excel is used to carry out the task.
b. After that, only the miRNAs associated with human and their expression values are chosen. In addition, we have removed every miRNA expression that had a null value. The resultant final data is used for tasks involving visualization, miRNA selection, and classification.

### iii. Data visualization

Two types of visualization plots are used in our study. All the classification methods are implemented using python packages.

a. To examine the data set’s five-number summary (minimum, first quartile, median, third quartile, and maximum) we used box and whisker plot; and
b. To check the distribution of the miRNA expressions associated with each class (i.e., normal and cancer), we use kernel density estimate (KDE) plot.

All results related to the data visualization are provided in Section 3.

### iv. miRNA selection

Prior to applying any classification model, it is necessary to select significant miRNAs that are associated with cancer development. A variety of methods, discussed below, are applied during the selection process and are executed by python packages.

a. Random Forest (RF) based selection: RF calculates the importance of a feature based on its capability to increase the pureness of the leaves. Higher the increment in leaves purity, by a feature, the higher the significance of it. The process is performed for every tree and the average importance is computed for all the trees. Finally, the importance value is normalized so that, the sum of the importance scores becomes 1.
b. Support Vector Machine (SVM) based selection: We fit the data to the linear SVM and accessed the classifier coefficients/weights of trained model. Feature importance is determined by comparing the value of these coefficients to each other. SVM coefficient is used to identify the main features for classification; thereby eliminating others from the selection process.
c. Extra Tree based (ET) selection: We first constructed a model with features. For each feature, the normalized total reduction in the Gini Index is computed to obtain a Gini importance value for that feature. To perform feature selection, each feature is ordered in descending order according to the Gini importance of each feature and we selected the top k features according to the selection criteria.
d. Chi-square information-based selection: Here we calculated the difference between expected count (E) and observed count (O). In other words, the relationship between the independent category (predictor) and the dependent category (response) is being determined here. For the feature selection purpose our goal is to choose the predictors (i.e., features) those are most relevant for the response (i.e., decision).
e. Maximum Relevance Minimum Redundancy-I (MRMR-I): It maximizes the relevance of a feature and minimizes the redundancy between multiple features. We computed mutual information between a feature and the corresponding class label for determining the relevance and the same operation between two different features for obtaining redundancy [16].
f. MRMR-II: Unlike MRMR-I, here instead of using mutual information-based theory, a classification-based model is used for computing the relevance.
g. Fuzzy Rough Entropy Measure (FREM): This measure emphasizes on entropy computation between miRNA expressions and class labels. The measure will give relevance of the miRNA. Note that lesser the value of entropy higher the relevance of a miRNA. Similarly, we computed the entropy between two miRNAs to find out redundancy between them. We selected less redundant miRNAs from the selected set of relevant miRNAs. Here the phenomena of class overlapping is handled by fuzzy set and the inexactness in the class sizes is dealt using rough lower approximation.

### v. Classification

After obtaining multiple sets of selected miRNAs, we used their (both identity and expression values) as features for classifying individuals having cancer from those who do not have cancer. We employed seven classifiers in this study, as briefly explained below. All the classification methods are implemented by python scripts.

a. RF: The random forest algorithm is special type of bagging method. It is made up of a group of decision trees, each of which has a subset of data taken from the entire training set with replacement. Bootstrap sampling is the process of drawing a subset of data for this technique.
b. SVM: It is a two-class classifier that is effective and operates by optimizing the hyperplane that separates the two classes. In case where data cannot be separated linearly, it is mapped to a high-dimensional feature space to determine perform the classification.
c. ET: This is an ensemble machine learning model (ML) that combines the output of several decision trees that have been trained to determine the outcome. Like random forests, it selects different variants of the training set using bagging to guarantee that decision trees differ enough from each other. However, Extra Trees trains decision trees on the complete dataset rather than a subset of data.
d. XGboost: This algorithm is specifically designed for structured and tabular data. XGBoost is an implementation of gradient boosted decision trees designed for speed and performance. XGBoost is an extreme gradient boost algorithm, i.e., it is an ML model that works with large and complex datasets. XGBoost is also an ensemble modelling like Random Forest or ET, however it uses boosting technique instead of bagging for combining the base ML model.
e. ANN: It is inspired by the human brain and developed as per the working methodology of neurons. This model consists of an input layer, one or multiple hidden layers, and an output layer. Each node (artificial neuron), connects to another node with a weight. If the output of any individual node is above a specified threshold value, only then the node will send the output to the next layer. Otherwise, it will remain inactive.
f. k-NN: The k-NN algorithm is a popular ML model which relies on the idea that similar data points tend to have similar class labels. It calculates the distance (e.g., Euclidean distance) between the unknown sample with all training samples. Then, it identifies the k shortest distant neighbors of the unknown sample based on the previously calculated distances. The major number of the class labels, available in the selected set of kneighbors, is considered as the predicted label of the unknown.
g. Naïve Bayes: A Naive Bayes classifier is a probabilistic ML model that is used for classification task. It uses Bayes theorem to predict the class label of an unknown sample.

## 3. Results

The Box and Whiskers plots of randomly chosen 50 representative miRNAs, corresponding to the three studies are plotted in Fig. 2 for representation purposes. Note that, these plots are created before using any selection algorithm.

**Fig. 2.**
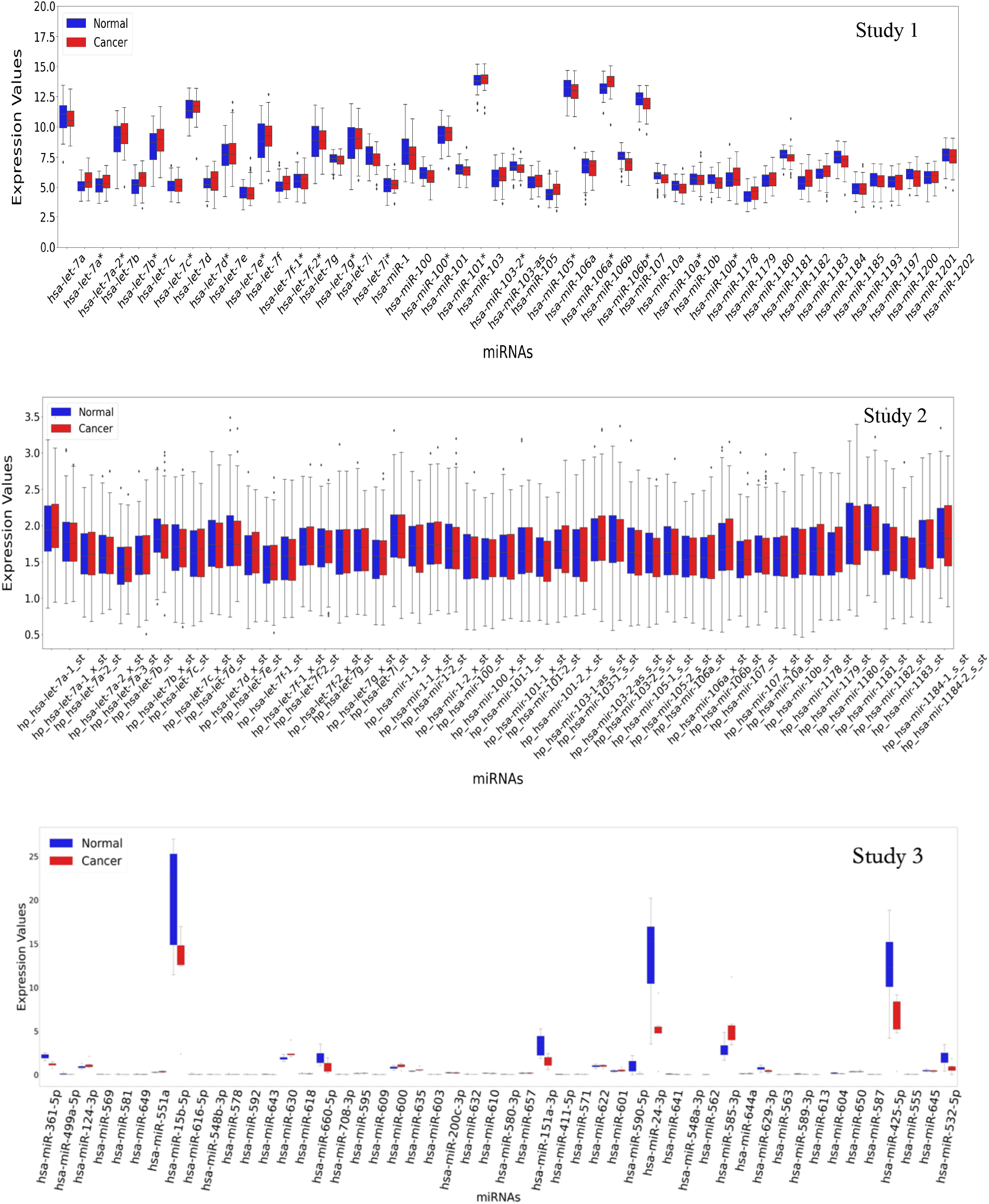
Box and Whisker plot for 50 representative miRNAs.

Furthermore, the estimated density (KDE plot) of all expressions is shown below in Fig. 3

**Fig. 3.**
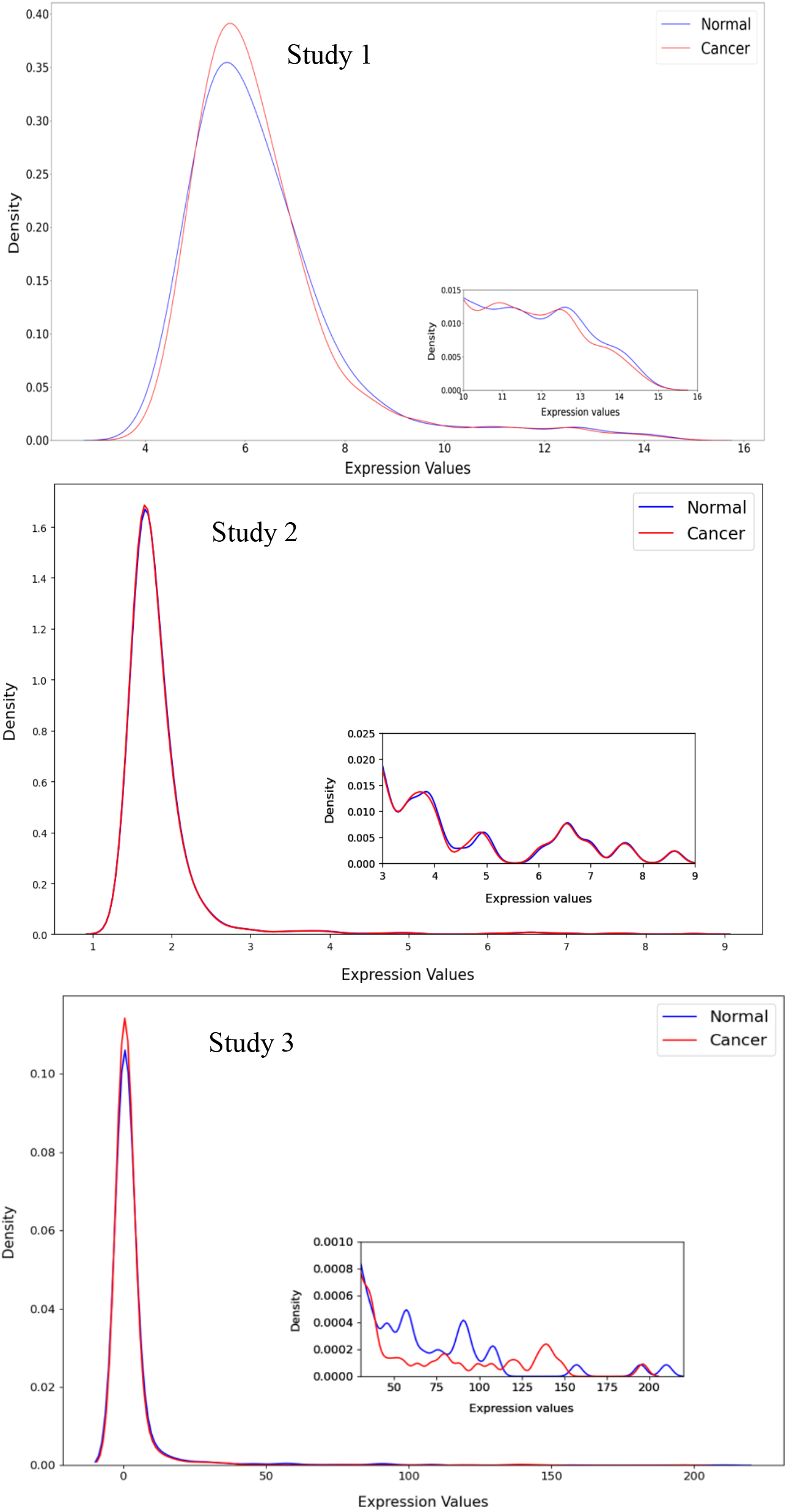
Density plot for normal and cancer expressions.

We applied different feature selection models to identify 10 most significant miRNAs from the entire data set, that correlate with cancerous samples. To assess the difference in the spread of expression values between cancer vs normal categories, we visualized the selected miRNAs using Box and Whisker plot. Fig. 4 shows the box plots of top 10 miRNAs using the best performing selection method corresponding to different studies.

**Fig. 4.**
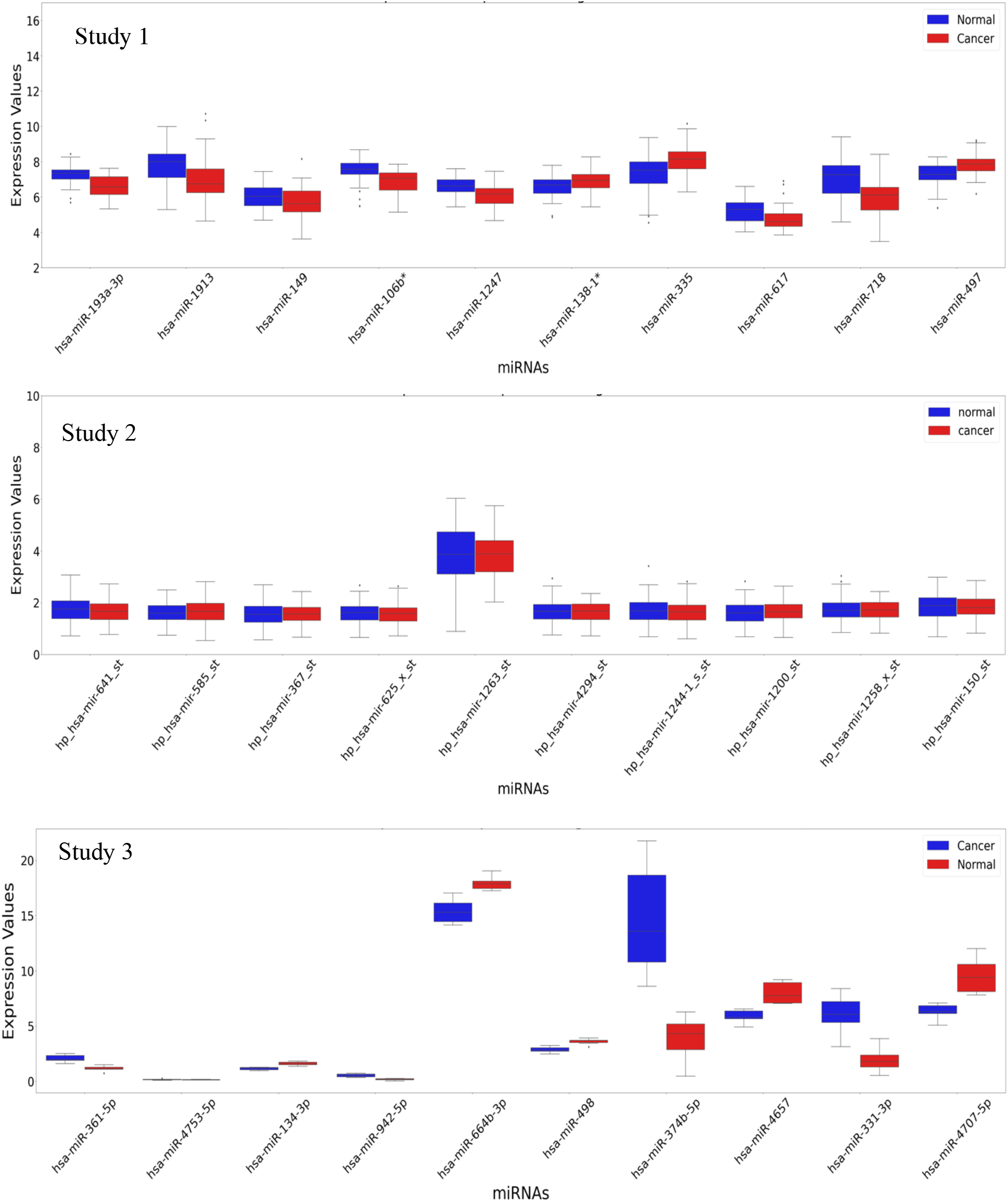
Top 10 significant miRNAs.

Table 2 reports the 10 best miRNAs corresponding to studies 1 & 3 obtained upon application of different feature selection models and can be considered as biomarkers for breast cancer diagnosis. “Best” refers to the output obtained from the selection algorithm that gives highest classification performance, in terms of accuracy. We observed from Fig. 4 that study 2 does not perform well in terms of class separation, hence no miRNAs from this study are shown in the Table 2.

**Table 2:** miRNAs shortlisted from Study 1 and 3 using MRMR-II algorithm.

| # | Study 1 | Study 3 |
| --- | --- | --- |
| 1 | hsa-miR-193a-3p | hsa-miR-361-5p |
| 2 | hsa-miR-1913 | hsa-miR-4753-5p |
| 3 | hsa-miR-149 | hsa-miR-134-3p |
| 4 | hsa-miR-106b* | hsa-miR-942-5p |
| 5 | hsa-miR-1247 | hsa-miR-664b-3p |
| 6 | hsa-miR-138-1* | hsa-miR-498 |
| 7 | hsa-miR-335 | hsa-miR-374b-5p |
| 8 | hsa-miR-617 | hsa-miR-4657 |
| 9 | hsa-miR-718 | hsa-miR-331-3p |
| 10 | hsa-miR-497 | hsa-miR-4707-5p |

Following the selection of miRNAs, we used multiple classifiers (section 6.v) to assess the potential of the selected miRNAs to be used as diagnostic biomarkers. Using these classifiers, we determined the accuracy, precision, recall, and F_1_-scores of the top 10 selected miRNAs. The graphs related to the accuracies of three different studies are shown in Fig. 5.

**Fig. 5.**
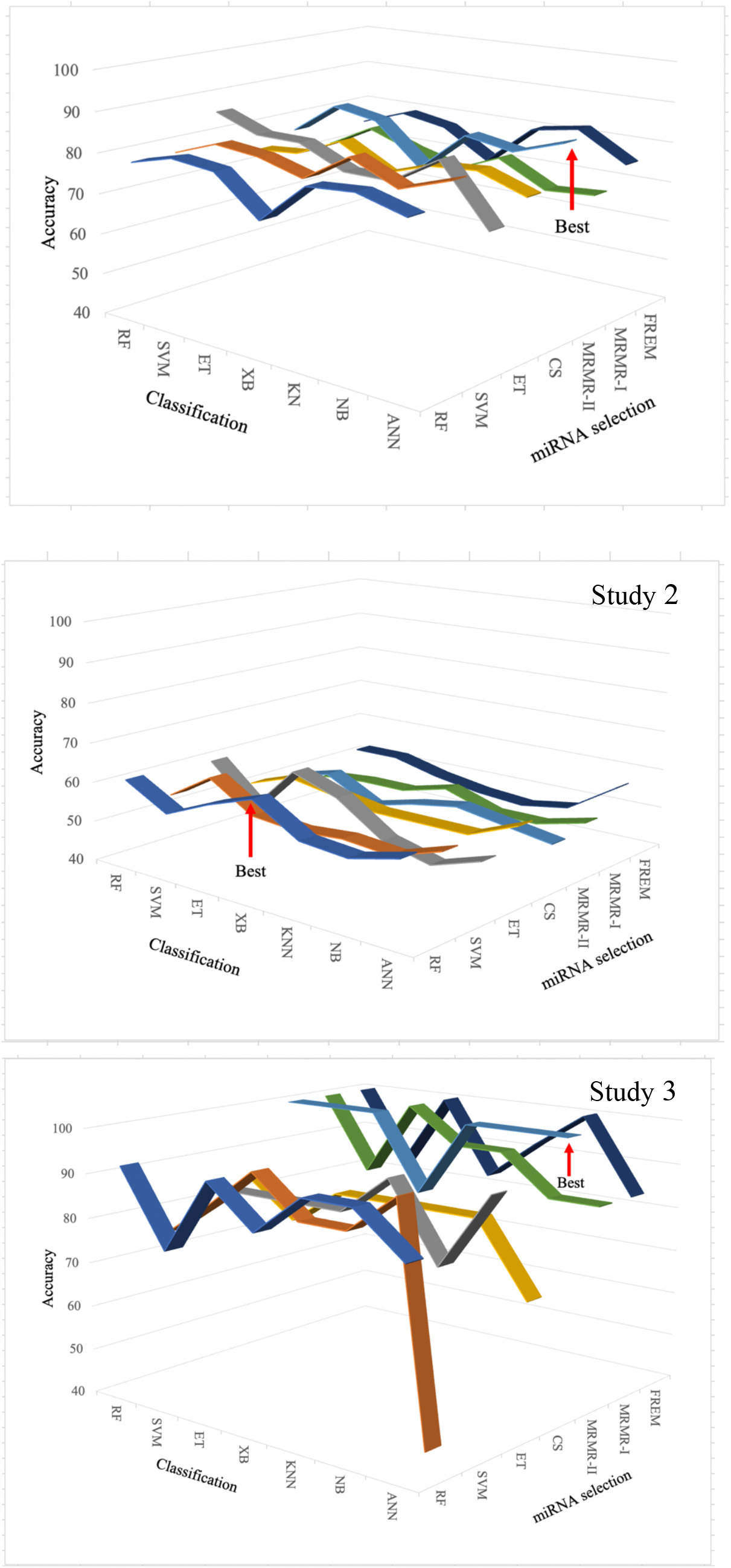
Comparison of selection-classification algorithm pairwise using top 10 miRNAs.

Additionally, we provide the confusion matrix corresponding to the best classification performance (using top 10 miRNAs) for these data sets, as shown in Fig. 6.

**Fig. 6.**
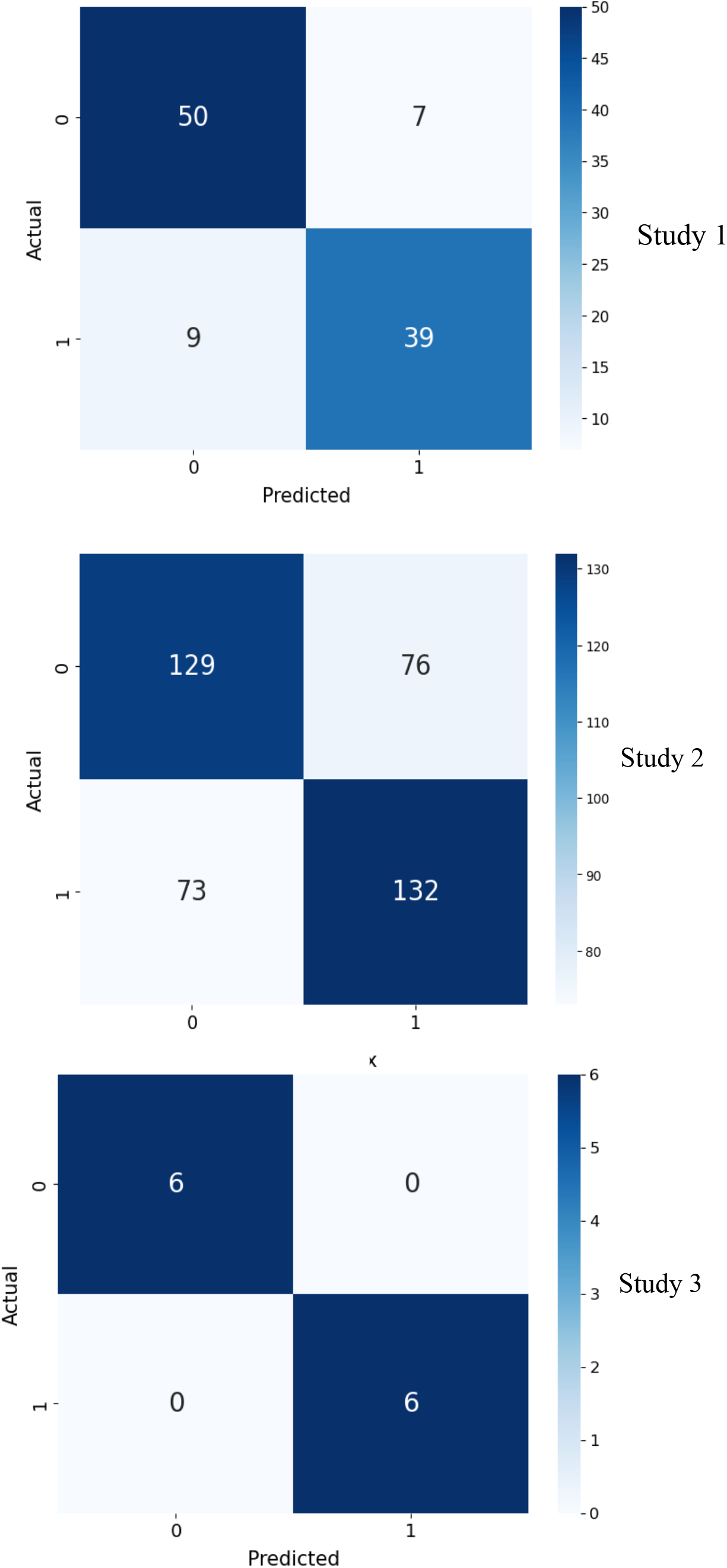
Confusion matrices.

Table 3 shows the results corresponding to the best-performing classification pairs

**Table 3:** Classification performance.

| Classifier | Precision<br>(Normal) | Precision<br>(Cancer) | Recall<br>(Normal) | Recall<br>(Cancer) | F1<br>(Normal) | F1<br>(Cancer) | Accuracy<br>(%) |
| --- | --- | --- | --- | --- | --- | --- | --- |
| **Study 1** |  |  |  |  |  |  |  |
| RF | 0.78 | 0.76 | 0.81 | 0.73 | 0.79 | 0.74 | 77 |
| SVM | 0.86 | 0.83 | 0.86 | 0.83 | 0.86 | 0.83 | 85 |
| ET | 0.83 | 0.83 | 0.86 | 0.79 | 0.84 | 0.81 | 83 |
| XGBoost | 0.75 | 0.72 | 0.77 | 0.69 | 0.76 | 0.70 | 73 |
| k-NN | 0.79 | 0.89 | 0.93 | 0.71 | 0.85 | 0.79 | 83 |
| Naïve Bayes | 0.86 | 0.76 | 0.77 | 0.85 | 0.81 | 0.80 | 81 |
| ANN | **<u>0.85</u>** | **<u>0.85</u>** | **<u>0.88</u>** | **<u>0.81</u>** | **<u>0.86</u>** | **<u>0.83</u>** | **<u>85</u>** |
| **Study 2** |  |  |  |  |  |  |  |
| RF | 0.61 | 0.58 | 0.63 | 0.60 | 0.60 | 0.62 | 61 |
| SVM | 0.55 | 0.55 | 0.55 | 0.56 | 0.55 | 0.56 | 55 |
| ET | 0.60 | 0.60 | 0.61 | 0.59 | 0.61 | 0.60 | 60 |
| XGBoost | **<u>0.64</u>** | **<u>0.63</u>** | **<u>0.63</u>** | **<u>0.64</u>** | **<u>0.63</u>** | **<u>0.64</u>** | **<u>64</u>** |
| k-NN | 0.56 | 0.58 | 0.63 | 0.50 | 0.59 | 0.54 | 57 |
| Naïve Bayes | 0.57 | 0.55 | 0.47 | 0.65 | 0.52 | 0.60 | 0.56 |
| ANN | 0.59 | 0.58 | 0.55 | 0.62 | 0.57 | 0.60 | 59 |
| **Study 3** |  |  |  |  |  |  |  |
| RF | 1.00 | 1.00 | 1.00 | 1.00 | 1.00 | 1.00 | 100 |
| SVM | 1.00 | 1.00 | 1.00 | 1.00 | 1.00 | 1.00 | 100 |
| ET | 1.00 | 1.00 | 1.00 | 1.00 | 1.00 | 1.00 | 100 |
| XGBoost | 0.83 | 0.83 | 0.83 | 0.83 | 0.83 | 0.83 | 83 |
| k-NN | 1.00 | 1.00 | 1.00 | 1.00 | 1.00 | 1.00 | 100 |
| Naïve Bayes | 1.00 | 1.00 | 1.00 | 1.00 | 1.00 | 1.00 | 100 |
| ANN | **<u>1.00</u>** | **<u>1.00</u>** | **<u>1.00</u>** | **<u>1.00</u>** | **<u>1.00</u>** | **<u>1.00</u>** | **<u>100</u>** |

## 4. Discussion

In this investigation we focus on blood sample-based breast cancer detection/diagnosis. This is a micro-invasive way of sample collection, and it facilitates a less stressful diagnostic procedure. While diagnosing with miRNAs enables early detection of cancer, the use of ML models speedup the whole procedure, making it doubly beneficial. These models can also reduce the cost required for biochemical experiments by providing the predictive results. However not all ML models are equally efficient for every problem or data sets. Therefore, we need to empirically check for the suitable ML models for this problem/task. Hence, in this study we extensively tested the available datasets on multiple well recognized selection methods and classifiers. It may be noted that our approach is applicable where the data related to both the classes (normal and cancer) are available. We have not considered any synthetic data generation procedure for our studies where one type of sample (normal / cancer) is not available. Another approach one can adopt; but not considered here, is where a classifier is trained with only a single category of data and given a test sample the model decides whether the sample belongs to the same category or not (one against all classifier).

The primary objective of this study is to identify the miRNAs that are associated with breast cancer and can serve as reliable biomarkers for cancer detection using a micro-invasive approach. It also investigates the significance of sample size and miRNA expression values in ML model-based diagnosis. Our datasets contain a wide range of patient counts, from relatively small (12 - 6 normal & 6 cancer) to medium (105 - 57 normal & 48 cancer) to large (410 - 205 normal & 205 cancer). For any data set with small number of samples (ex: study 1), it is difficult to maintain a separate validation data set apart from training & test set. This can lead to double dipping issues. “Double-dipping” (or, Data leakage) refers to a situation when both feature selection and classification are performed on the same samples using the same criteria. This can compromise the generalization capability of the model, and a reduction in the statistical significance of observed performance. This can in turn lead to misleading results and/or overfitting. In the case of overfitting, the model may produce good performance with the test data (since it was trained on the same) but with the unknown, real-world data the model may fail to perform.

Our study is not affected by double dipping as we perform feature selection in a separate step using multiple criteria (Models). As we are not focused on evaluating the performance of the classifiers a separate training set was not defined before the miRNA selection (i.e., the training and test set are not generated before miRNA selection, rather they are created after miRNA selection and before the classification process), because using this approach the selected set of miRNAs may change for each round of cross validation. If the selected miRNAs for a particular data set change at each stage of cross validation, then there will be no generalization of selected miRNAs. This is not a desirable situation in this study, because the main goal is the identification of miRNAs that correlate/associate with breast cancer. Furthermore, the importance of a set of miRNAs can only be determined if the classification model uses a fixed set during cross validation, rather than a different set each time. Similar approach for miRNA selection can be observed in [18]. To maintain uniformity we followed the same procedure for comparatively medium and large data set also.

After selecting the miRNAs, we attempt to classify the samples (i.e., patients) as normal or cancer, using miRNA’s as features (using their expression values). We achieve this by the training & testing of multiple classification models on the selected miRNA’s. This ensures that the classification models are getting the scope to perform with the features selected by different algorithms (as compared to the algorithm used in the selection step). This is one of the approaches to avoid double dipping [19]. To evaluate the performance of these miRNA’s (reflected in Table 3) and to improve the generalization, we employ a widely used 5-fold cross validation approach. In this approach the total number of samples are divided into 5 equal (or approximately equal) subsets. For each round of training/testing of a classification model, one subset is used as the test set and the remaining four are used as training set. This process is repeated (i.e., five rounds) by considering each subset as the test set and the remaining ones as training. The average results obtained from the said five rounds are considered as the results of the corresponding classification model.

The novelty of our research article lies in combining three aspects. The first aspect is the micro-invasive sampling-based approach (blood) to detect miRNA to diagnose breast cancer. The micro-invasive procedure to collect samples ensures diagnosis with minimal pain. The second aspect is the use of different ML based feature selection algorithms to identify the miRNAs associated with breast cancer. The use of multiple models/criteria for miRNA selection reduces the chance of overfitting while application of the classifiers. The final aspect is evaluating the performances of these selected miRNAs with various classifiers; thereby observing the robustness of these ML models in feature selection and classification.

Fig. 2 demonstrates that it is not possible to directly distinguish visually between miRNA expression patterns of normal vs cancer cases. Hence, there is a need to select those miRNA’s (amongst the thousands) that are significantly associated with the cancer development. In this regard, we selected a set of 10 top miRNAs (Table 1). Since the selected miRNAs in study 2 do not exhibit any major differentiation between normal and cancer groups (Fig. 3, study 2) hence in Table 1, we reported only two studies (i.e., 1 & 3). This table also shows that across the two shortlisted datasets there are no common miRNAs. Since the patient data sets originated from different countries (Germany, & China) across different continents, it is possible that the set of significant miRNAs can vary based on the population genetics across different geographic locations. This suggests that location-based, population-type specific identification of cancer biomarkers will be key to a tailored, precision bio-marker strategy.

Fig. 4 demonstrates that in studies 1 & 3 there are clear differences between the expression patterns of miRNAs in normal vs cancerous conditions. However, for study 2 the difference is not apparent even after using top 10 miRNAs. The observations of studies 1 & 3 indicate the success of blood sample-based breast cancer diagnosis strategy employing miRNA expression values. Additionally, from the density estimation plot (Fig. 3) we observed that miRNA expression in both the studies follow Gaussian distribution. In Study 1 (Table 1) there are 57 normal subjects and 48 cancer patients. Fig. 3 (Study 1) shows that the expression range varies from 3-16, and Fig. 4 (Study 1) shows that most of the selected miRNAs have higher values of expressions (between 6 – 10). In Study 2 (Table 1), there are 205 cancer patients and normal subjects each. However, it is observed from the corresponding density estimation plot (Fig. 3, Study 2) that majority of the expression values are ≤ 4. In the selected set of miRNAs (Fig. 4, Study 2), we observed that both the categories (normal & cancer) do not show much difference in terms of class spread and overlap with each other. Each category in Study 3 (Table 1) contains only six individuals. Here it is observed that the expression values are higher than 20 (Fig 3, Study 3). Interestingly, after selecting the top 10 miRNAs (Fig. 4, Study 3) we observe large difference in terms of expression spreads in both the categories which is also reflected in the classification results.

Upon careful observation of the results corresponding to these three data sets, we believe that the sample size has a relatively minor influence in the selection/classification of normal vs cancer. Table 3 shows that Study 1 produces good results, with a moderate sample size (mentioned in Table 1). The classification performance of Study 2 gets significantly low as seen in Table 3, despite having comparatively large number of samples in each category (mentioned in Table 1). This reflects that low expression levels of miRNA are affecting the detection/diagnosis irrespective of sample size. This claim gets support from the results of Study 3 where we have small number of samples (mentioned in Table 1), but high miRNA expression values. In this case, superior performance in terms of classification is observed in Table 3 compared to the other data sets. The claim regarding the sample sizes gets further support from the confusion matrices (Fig. 6). Here, we observe that the number of true positive and false positives are not dependent on the sample size, rather it is the level of miRNA expression values that are critical. Our realization also gets support form Fig. 4 where we can see that the majority of the selected miRNAs belong to the higher range of expression values in the density plot (see Fig. 3).

In terms of classification performance, it is observed (Fig. 5) from the results that the MRMR-II provides the best set of miRNAs for the two most successful data sets of study 1 & 3. The random forest-based selection provides the best result for study 2. From the confusion matrices (Fig. 6, Study 2) we get the indication of low precision/recall/F_1_-score/accuracy. Despite having a large sample size in Study 2, the corresponding confusion matrix with the best results shows that the data set produces many false positives/negatives due to the low magnitude of the miRNA expression values. On the other hand, for studies 1 & 3 (Fig. 6) we obtained satisfactory numbers of true positives/negatives due to the availability of comparatively high miRNA expression values. In Table 3, it is observed that the precision, recall, F_1_ and accuracy values are 0.8-1.0 / 80-100% for Studies 1 & 3, and these are ∼0.60/60% for Study 2. In this table the row of the best performing classifier is marked by bold and underlined fonts.

For both the studies 1 and 3 it is observed that information theory-based miRNA selection procedure (MRMR-II) is most successful. For Study 2 a classifier (RF) based selection method claimed the best position. It may be noted that in all the data sets the number of samples (patients/normal people) are much less compared to the high number of miRNAs (features). Given the circumstances where the sample size is even smaller (Studies 1 & 3), it is unlikely to get good performance with traditional classifier-based feature selection. Accordingly, we observe better performance for studies 1 & 3 using MRMR-II feature selection method. On the other hand, RF-based selection method provided better performance for study 2 as it has larger set of samples. However, it may be noted that, with study 2 we were unable to obtain satisfactory results in classification.

In our study, ANN provides the best classification results for Study 1 and 3. It is a promising classifier, and it works based on assigning weights to the features for the classification. It can be constructed by one or multiple hidden layers to capture the intrinsic data patterns; thereby producing superior performances. The data visualization of study 3 (Fig. 4) clearly demonstrates that the range of expression distributions, corresponding to the normal and cancer samples, are different. This can be considered as the likely reason for success with the classifiers. As expected, in classification phase we obtained superior results, and all classifiers (except XGBoost) produced 100% performance in terms of various measures (Table 3).

Our approach essentially requires two different categories/classes to detect/diagnose cancer. It is possible that in some publicly available datasets, only the cancer patients and their corresponding miRNA expression values are available, whereas the normal subject data is not available. In such cases our approach will have limited application, as in this study, we have not emphasized on any synthetic data generation technique or didn’t view the task as a one against all classification problem. These will form the basis of our future investigations.

## Conclusions

Our analysis highlights multiple advancements of blood-based breast cancer diagnosis approach. First, it shows that modest-high expression levels of miRNA are required to be presented to any ML model. This is also reflected in the density plots of miRNA expressions, where it is seen that all the selected miRNAs occupy the high expression range in the gaussian plot. Second, we show that the dysregulation pattern of selected miRNAs can reflect both an up/down regulation under cancerous condition, which could hint at their biological role as suppressors/enhancers of the disease phenotype. The evaluation result shows the promising performance using the selected set of miRNAs. Therefore, we conclude that if the miRNA expression values have a good magnitude, then they can be used as effective biomarkers for blood-based breast cancer diagnosis.

## Author contributions

JKP and BR conceived the study. JKP carried out the analysis. JKP and BR created the manuscript framework and wrote the manuscript. JKP and BR interpreted the results. BR supervised the study, edited the manuscript for scientific and language accuracy, finetuned the framework and interpretations; provided critical suggestions.

## Acknowledgements

JKP and BR acknowledge Jio Platforms Limited, for providing the necessary facilities required and financial assistance through the salaries provided.

